# Hypoxic Preconditioning Promotes Survivals of Human Adipocyte Mesenchymal Stem Cell via Expression of Prosurvival and Proangiogenic Biomarkers

**DOI:** 10.1101/2021.01.18.427057

**Authors:** I Gde Rurus Suryawan, Budi Susetyo Pikir, Fedik Abdul Rantam, Anudya Kartika Ratri, Ricardo Adrian Nugraha

## Abstract

**Background:** Contributing factors for improved survival of human adipocytes mesenchymal stem cells (h-AMSCs) cultured through hypoxia preconditioning, in example apoptosis inhibition involving BCL2 and HSP27 expression, trigger signal expression (VEGF), SCF expression, OCT-4 expression, and CD44+ expression.

**Objective:** To explain the mechanism and role of hypoxic preconditioning and the optimal duration of hypoxic preconditioning exposure to improve survival of h-AMSCs so that could it could be used as a benchmark for h-AMSCs culture strategy before transplantation.

**Methods:** This study was an experimental laboratory explorative study (in vitro study) with hypoxic preconditioning in human-adipose mesenchymal stem cells (h-AMSCs) cultures. This research was conducted through 4 stages ;First, Isolation and h-AMSCs culture from adipose tissue of patient (human). Second is the characterization of h-AMSCs from adipose tissue by phenotype (Flowcytometry) through CD44+, CD90+ and CD45-expression before being pre-conditioned for hypoxic treatment. Third, the hypoxic preconditioning in h-AMSCs culture (in vitro) was performed with an oxygen concentration of 1% for 24, 48 and 72 hours. Fourth, observation of survival from h-AMSCs culture was tested on the role of CD44 +, VEGF, SCF, OCT-4, BCL2, HSP27 with Flowcytometry method and apoptotic inhibition by Tunnel Assay method.

**Results:** The result of regression test showed that time difference had an effect on VEGF expression (*p*=0,000; **β**=−0,482) and hypoxia condition also influenced VEGF expression (*p*= 0,000; **β**=0,774). The result of path analysis showed that SCF had an effect on OCT-4 expression (*p*=0,000; **β**=0,985). The regression test results showed that time effects on HSP27 expression (*p*=0.000; **β**=0.398) and hypoxia precondition also affects HSP27 expression (*p*=0.000; **β**=0.847). Pathway analysis showed that BCL2 expression inhibited apoptosis (*p*=0.030; **β**=−0.442) and HSP27 expression also inhibited apoptosis (*p*=0,000; **β**=−0.487).

**Conclusion:** In conclusion, hypoxic preconditioning of h-AMSC culture has proven to increase the expression of VEGF, SCF, OCT-4, and BCL2 and HSP27. This study demonstrated and explained the existence of a new mechanism of increased h-AMSC survival in cultures with hypoxic preconditioning (O2 1%) via VEGF, SCF, OCT-4, BCL2, and HSP 27. But CD 44+ did not play a role in the mechanism of survival improvement of human AMSC survival.

## Introduction

Several literatures provide abundant information that human adipocytes mesenchymal stem cells (h-AMSCs) is an attractive resource for therapeutics and have beneficial effects in regenerating injured cardiomyocytes due to their self-renewal ability and broad differentiation potential under physiological and pathological conditions.^1–3^

Despite the impressive potential of the h-AMSC-based therapy, several obstacles (e.g., the difficulty of maintaining self-renewal and poor survival due to apoptosis and/or necrosis at the administration site) have been encountered.^4^ Some studies suggest that more than 90% of transplanted stem cells, either intravenously, intramyocardially, and intracoronary delivery, have necrosis and apoptosis and only about 5% transplanted stem cells can survive up to 14 days in infarcted myocardium.^5^ The survival of stem cells transplantation is so poor because high percentage of dead cells due to factors such as limited availability of blood, hypoxia, oxidative stress, inflammatory processes, loss of extracellular cell buffer matrix (anoic), non-conducive microenvironment to myocardial infarction, structural damage to blood vessels and lack of nutritional support.^6^

Therefore a particular strategy is needed to improve survival, increase proliferation, migration, maintain the potential for differentiation and viability of stem cells in environments with low oxygen levels. One of those strategies is to pre-condition hypoxic precursors in vitro on oxygen concentrations mimicking the stem cells’ niche.^7,8^ Contributing factors for improved survival of h-AMSCs cultured through hypoxia preconditioning, i.e., apoptosis inhibition involving BCL2 and HSP27 expression, trigger signal expression (VEGF), SCF expression, OCT-4 expression, and CD44 + expression.^9^

In detail, it has never been explained how far the role of hypoxic preconditions in inhibiting apoptosis of h-AMSCs culture in vitro, in order to enhance survival and increase proliferation, maintain multi-potency, stemness and inhibition of apoptosis. Based on the description above, we consider it is necessary to conduct a research to explain the increased survival of h-AMSCs through the treatment of sub-lethal hypoxia precondition (oxygen concentration of 1%) for 24, 48, and 72 hours by looking at the expression of inhibition on apoptosis and HSP27 expression, and BCL2. In addition, it is necessary to observe the role of hypoxic preconditions in the proliferation process through the expression of SCF, OCT-4, and BCL2.

## Objective

A study was conducted to explain and confirm the mechanism and role of hypoxic preconditioning and the optimal duration of hypoxic preconditioning exposure to improve survival of h-AMSCs so that could it could be used as a benchmark for h-AMSCs culture strategy before transplantation. This study was an experimental laboratory explorative study (in vitro study) with hypoxic preconditioning in human-adipose mesenchymal stem cells (h-AMSCs) cultures.

## Methods

### Ethical approval

The use of humans subjects in this study had been obtaining an ethical approval from research ethics committee of Dr. Soetomo Academic General Hospital - Faculty of Medicine, Airlangga University (Number: 264/Panke.KKE/IV/2017) issued on April 6^th^, 2017 under the name of I Gde Rurus Suryawan as principal investigator.

### Study design

This study is an exploratory laboratory experimental study (in-vitro study) with hypoxic preconditions in the culture of human-adipose mesenchymal stem cells (h-AMSCs) derived from human adipose tissue. The aim of this study was to obtain stem cells that have high survival so that they are not only viable but also have high adaptability to the environment when the stem cells are transplanted. This type of experiment is a true experimental post-test only control group design accompanied by phenotypic h-AMSCs characterization against CD44+, CD90 + and CD45-before being given treatment.

### Study setting

This research was conducted at the Center for Research and Development of Stem Cell-Institute Tropical Disease (ITD) Universitas Airlangga, Dr. Soetomo Academic General Hospital and the Faculty of Medicine, Airlangga University, Surabaya. The implementation of this study lasted for 2-3 months.

### Sample size

The sample size in this study was obtained using the Federer’s formula for sample size.^10^ This formula is used as a control for the degree of freedom in MANOVA. The formula description is as follows:

Sample size: (r-1) (K-1) ≥ 15
r = replication (experimental unit sample size per group)
K = number of subject group observations
K = 6 □ (r-1) (K-1) ≥ 15
(r-1) (6-1) ≥ 15
(r-1) 5 ≥ 15 □ r-1 ≥ 3 □ r = 4

Then the number of replications for each group is 4, so that the total sample is 24 plate culture.

### Materials

Experimental Unit:

1. h-AMSCs, namely human-adipose mesenchymal stem cells from adipose tissue obtained from minimally invasive surgery with small incisions (3-5 cm) under local anaesthesia in the lower abdominal area by a surgeon (*Figure 1*). These patients were patients who were prepared for clinical application of stem cell therapy by the Network Bank team at Dr. Soetomo Hospital, Surabaya. After getting the consent (the informed consent for fat tissue was taken) and we made separate informed consent about the use of some fat tissue for our study. The h-AMSCs experimental unit was taken from adult patients who were in a stable state who were not taking anti-platelets or anti-coagulants and then multiplied in vitro at the 5^th^ passage to 24 units. A total of 24 units were divided into 2 groups, namely control and treatment. The control group (P0) had 12 culture units in normoxic conditions (21% O_2_ concentration). The treatment group (P1) was 12 units pre-conditioned to hypoxia (1% O_2_ concentration). Both treatment groups were observed for survival (CD44+, VEGF, SCF, OCT-4, BCL2, HSP27, and apoptotic inhibition at 24, 48 and 72 hours of cell culture). Observation of apoptotic inhibition based on the expression of BCL2 and HSP27 along with the percentage of apoptosis that occurred.
2. Washing buffer (phosphate-buffered saline, PBS, Sigma-Aldrich, Milan, Italy, 0.1% sodium azide, and 0.5% bovine serum albumin (BSA), Radnor, USA) was used for all washing steps (3 ml of washing buffer and centrifugation, 400 g 8 min at 4°C). Briefly, 5×10^5^ cells/sample were incubated with 100 ml of 20 mM ethylenediaminetetraacetic acid (EDTA, Sigma-Aldrich) at 37°C for 10 min and washed.

**Figure 1.**
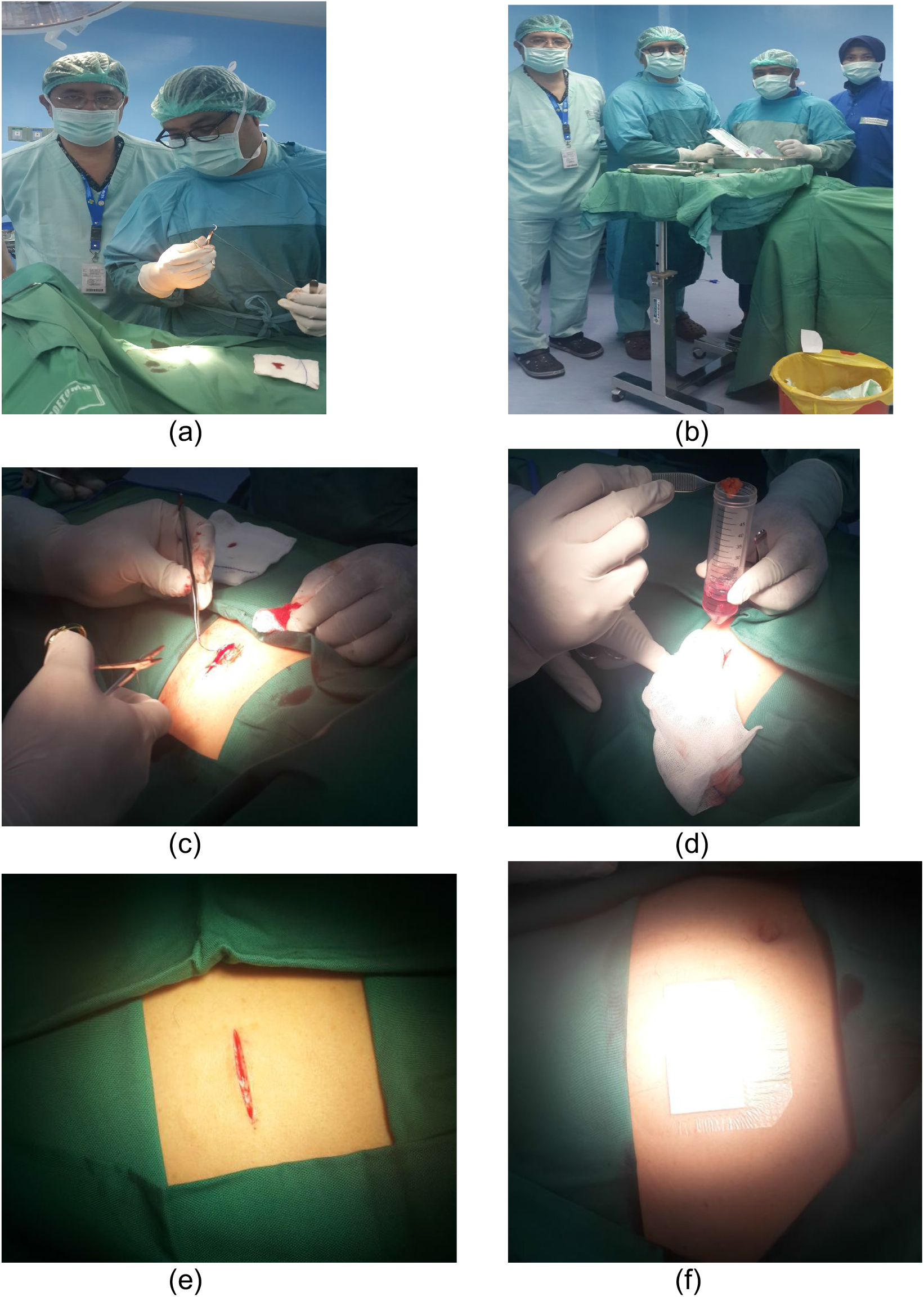
Isolation and culture of h-AMSCs from the patient’s adipose tissue (human)

### Experimental procedures

This research was conducted in 4 stages as follows:

1. Isolation and culture of h-AMSCs from the patient’s adipose tissue (human) (F*igure 1*)
2. Characterization of h-AMSCs from adipose tissue phenotypically (Flowcytometry) through identification of CD44+, CD90+ and CD45-before being treated with hypoxic preconditions.
3. Hypoxic precondition in in vitro h-AMSCs culture was carried out with an oxygen concentration of 1% for 24, 48 and 72 hours.
4. Observation of survival of h-AMSCs in the form of CD44 +, VEGF, SCF, OCT-4, BCL2, HSP27 expression, and apoptotic inhibition:

a. Phenotype expression of CD44 + was carried out by the flowcytometric method.
b. Immuno-cytochemical expression of VEGF
c. Immunocytochemical expression of SCF from h-AMSCs culture
d. Phenotype of OCT-4 expression (Immunocytochemistry and Immunofluorescence)
e. Apoptotic inhibition, based on the expression of BCL2 and HSP27 by immunocytochemistry accompanied by a low percentage of apoptosis through the Tunnel Assay method (*Figure 2*).

**Figure 2.**
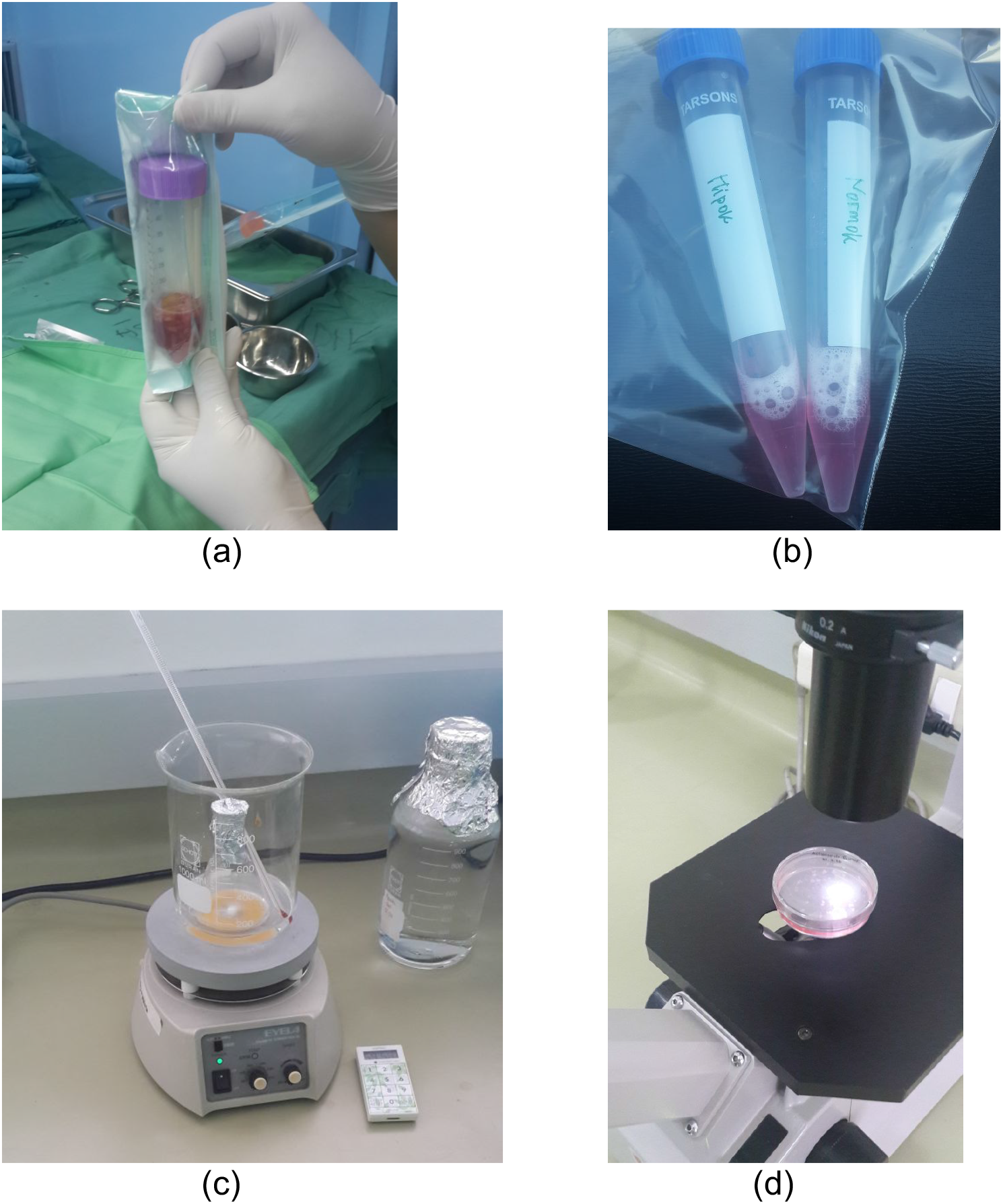
Observation of survival of h-AMSCs in the form of CD44 +, VEGF, SCF, OCT-4, BCL2, HSP27 expression, and apoptotic inhibition: a. Phenotype expression of CD44 + was carried out by the flowcytometric method. b. Immuno-cytochemical expression of VEGF c. Immunocytochemical expression of SCF from h-AMSCs culture d. Phenotype of OCT-4 expression (Immunocytochemistry and Immunofluorescence)

### Data analysis

Data collected, processed and statistically tested with several stages. The first stage is an Assumption Test in the form of a normality test to ensure that the data is normally distributed. Furthermore, a comparison test was carried out between the treatment group and the control group using Multivariate Analysis of Variance (MANOVA). Furthermore, path analysis is carried out to determine the pathway mechanism of the influence of the independent variables on the dependent variable by using multiple linear regression statistical tests. The statistical analysis was used to explain the effect of time (24, 48 and 72 hours) and hypoxic conditions on the expression of VEGF, SCF, OCT-4, CD44 +, BCL2, HSP27 and the number of cells undergoing apoptosis. The data scale of each variable under study is a ratio, so it is appropriate to use the Multiple Linear Regression statistical test. Statistical tests were performed using SPSS version 24.0 software.

## Results

The results showed that the time difference test on CD44+ expression was 24 hours with 48 hours (*p* = 0.017), 24 hours with 72 hours (*p=*0.004), and 48 hours with 72 hours (*p*=0.801). The result of regression test showed that time difference had an effect on expression of CD44 + (*p* = 0.002, **β**= −0.582) and hypoxia condition had no effect to CD44 + expression (*p* = 0.066, **β**= 0,317) (*Table 1*) (*Figure 3*).

**Figure 3.**
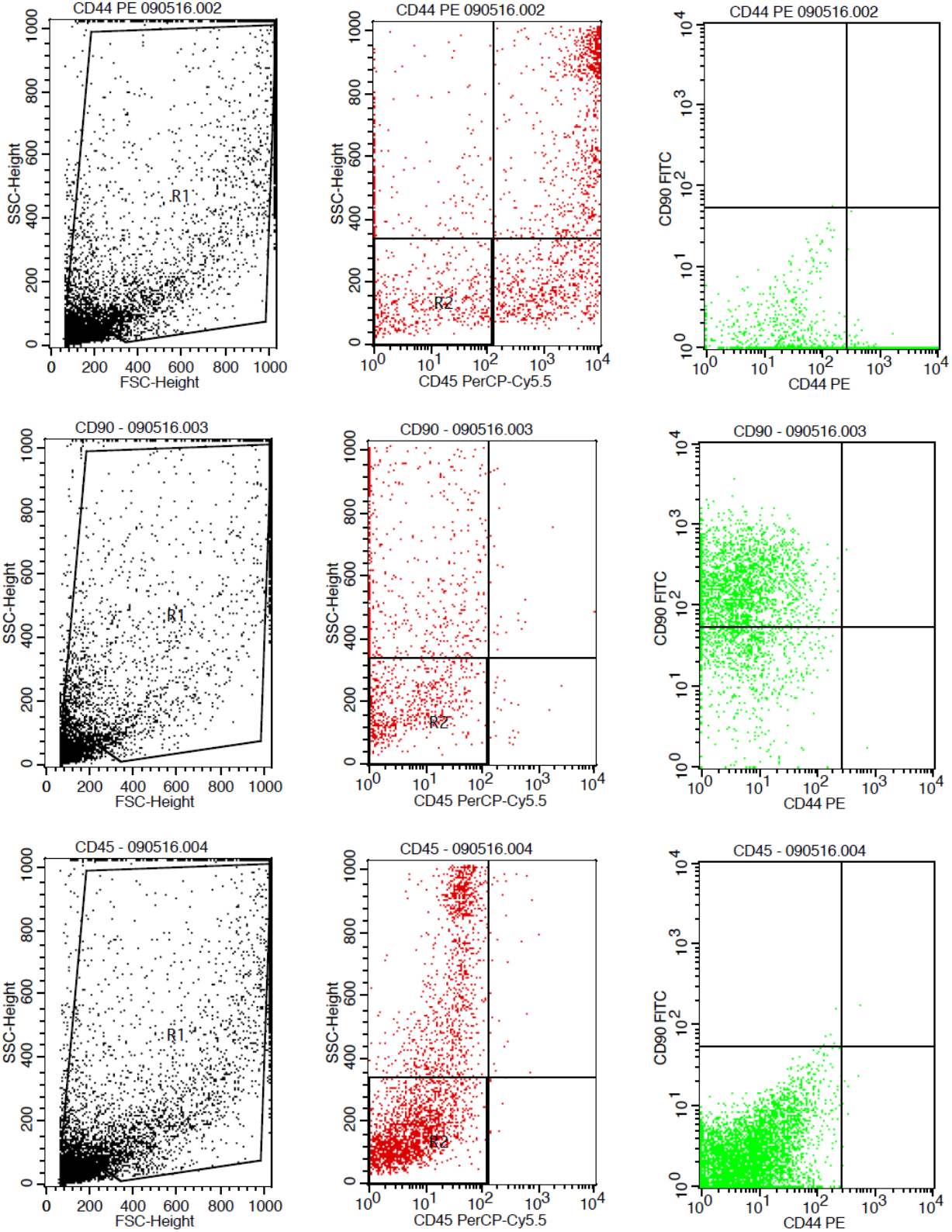
Flowcytometry results from human AMSCs based on cell culture for CD44+ CD90+ CD45-expression

**Table 1.**
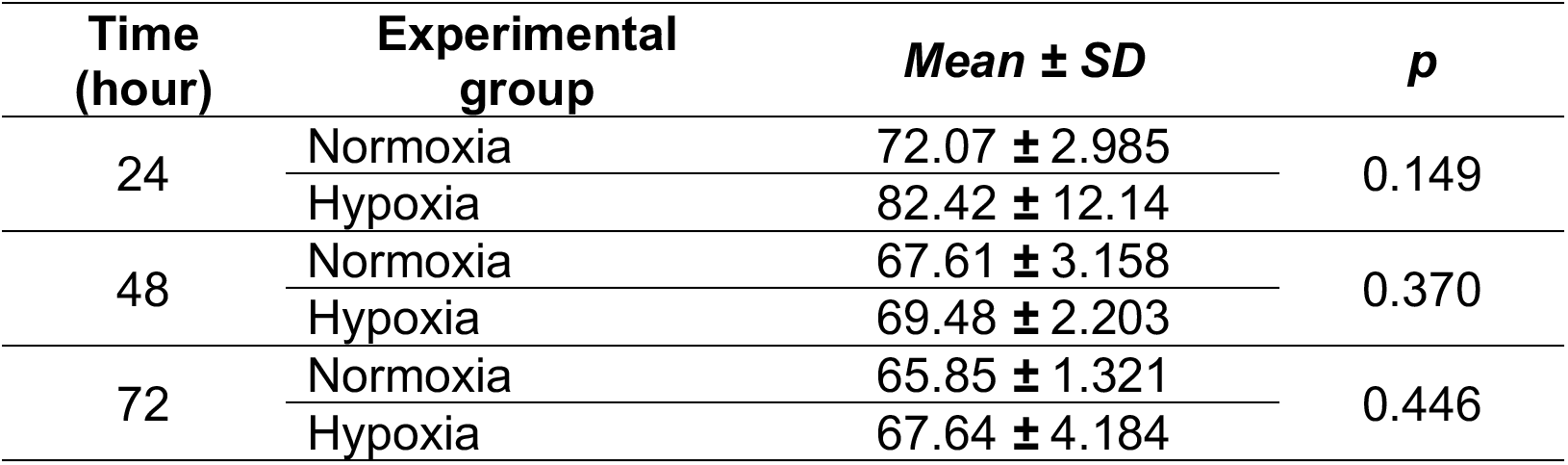
Results on CD44+ expression

| Time (hour) | Experimental group | Mean ± SD | p |
| --- | --- | --- | --- |
| 24 | Normoxia | 72.07 ± 2.985 | 0.149 |
| | Hypoxia | 82.42 ± 12.14 | |
| 48 | Normoxia | 67.61 ± 3.158 | 0.370 |
| | Hypoxia | 69.48 ± 2.203 | |
| 72 | Normoxia | 65.85 ± 1.321 | 0.446 |
| | Hypoxia | 67.64 ± 4.184 | |

The result of time difference test on VEGF expression is between 24 hours with 48 hours (*p*< 0.001), 24 hours with 72 hours (*p*<0.001), and 48 hours with 72 hours (*p=*0.047). The result of regression test showed that time difference had an effect on VEGF expression (*p*<0.001; **β**=−0,482) and hypoxia condition also influenced VEGF expression (*p*<0.001; **β**= 0,774) (*Table 2*) (*Figure 4*).

**Figure 4.**
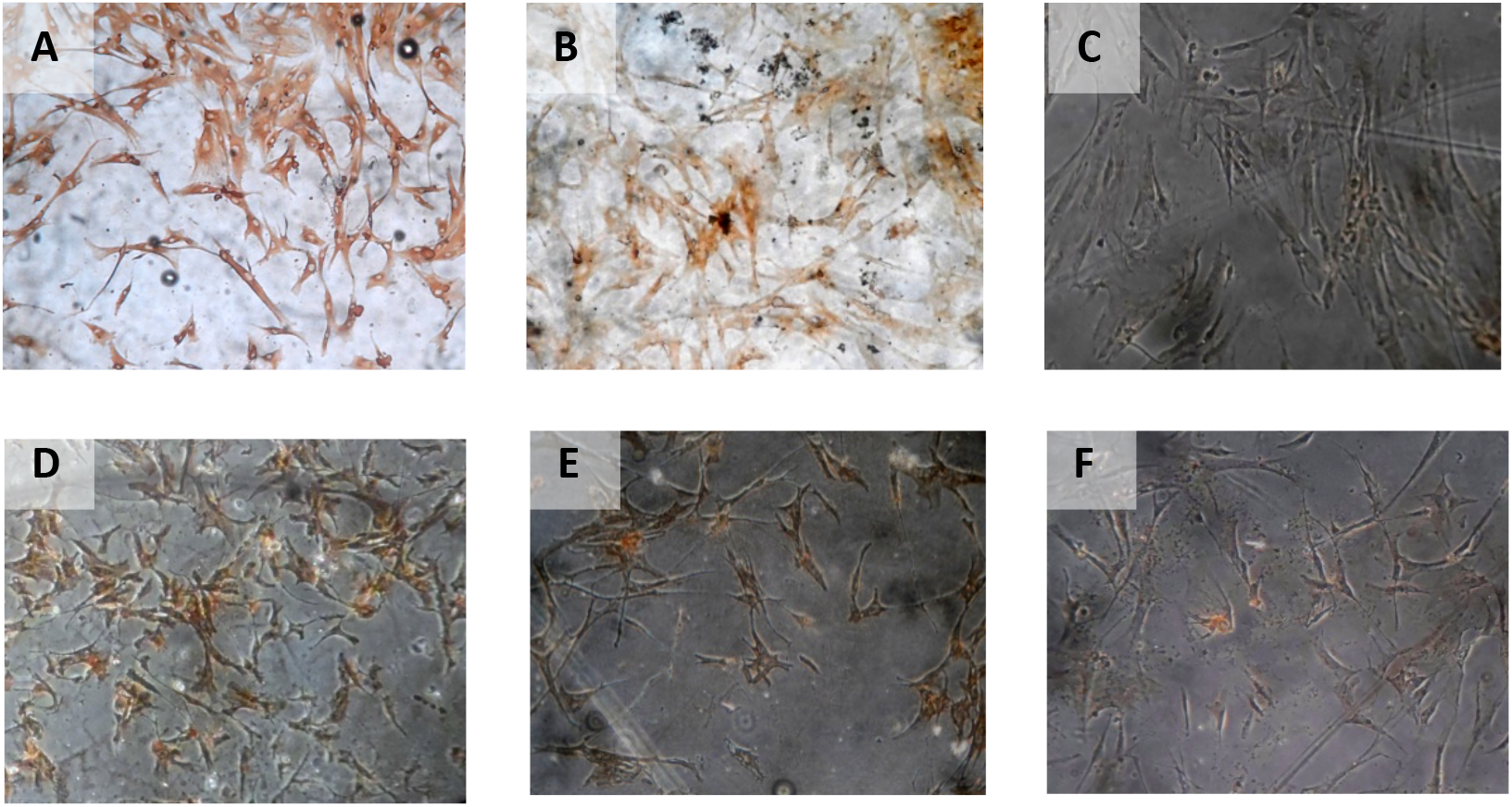
Immunohistochemical Characteristic of *h-AMSCs* based on VEGF expression at: a) normoxic condition for 24 hours; b) normoxic condition for 48 hours; c) normoxic condition for 72 hours; d) hypoxic condition for 24 hours; e) hypoxic condition for 48 hours; f) hypoxic condition for 72 hours.

**Table 2.**
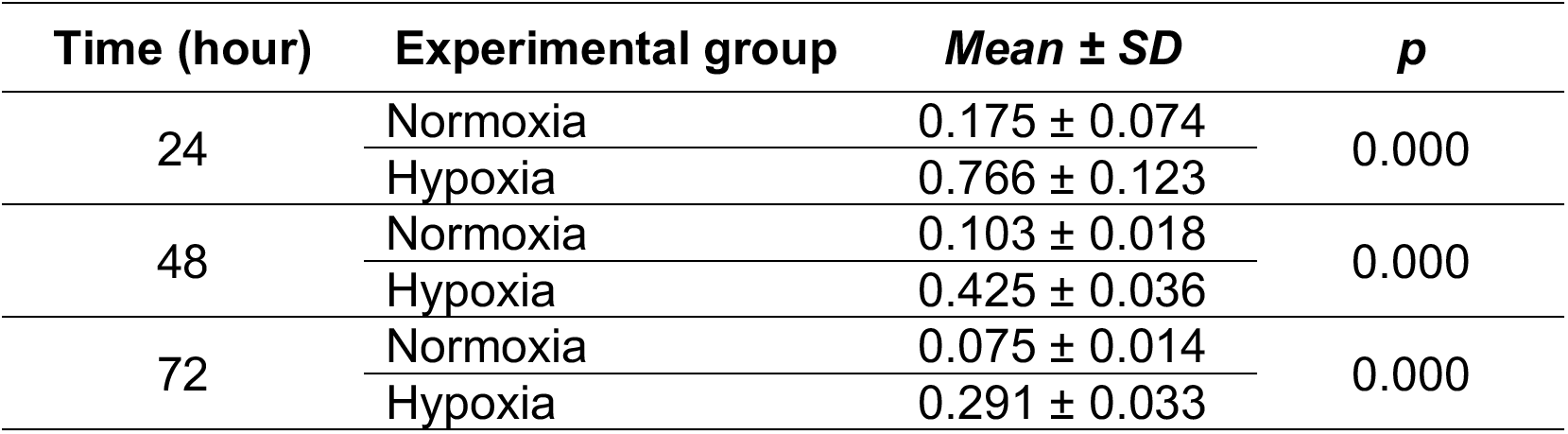
Results on VEGF expression

The result of time difference test on SCF expression is between 24 hours with 48 hours (*p*=0.283), 24 hours with 72 hours (*p*<0.001), and 48 hours with 72 hours (*p*<0.001). The result of path analysis showed that VEGF influenced the expression of SCF (*p*<0.001; **β** =0.889). (*Table 3*) (*Figure 5*).

**Figure 5.**
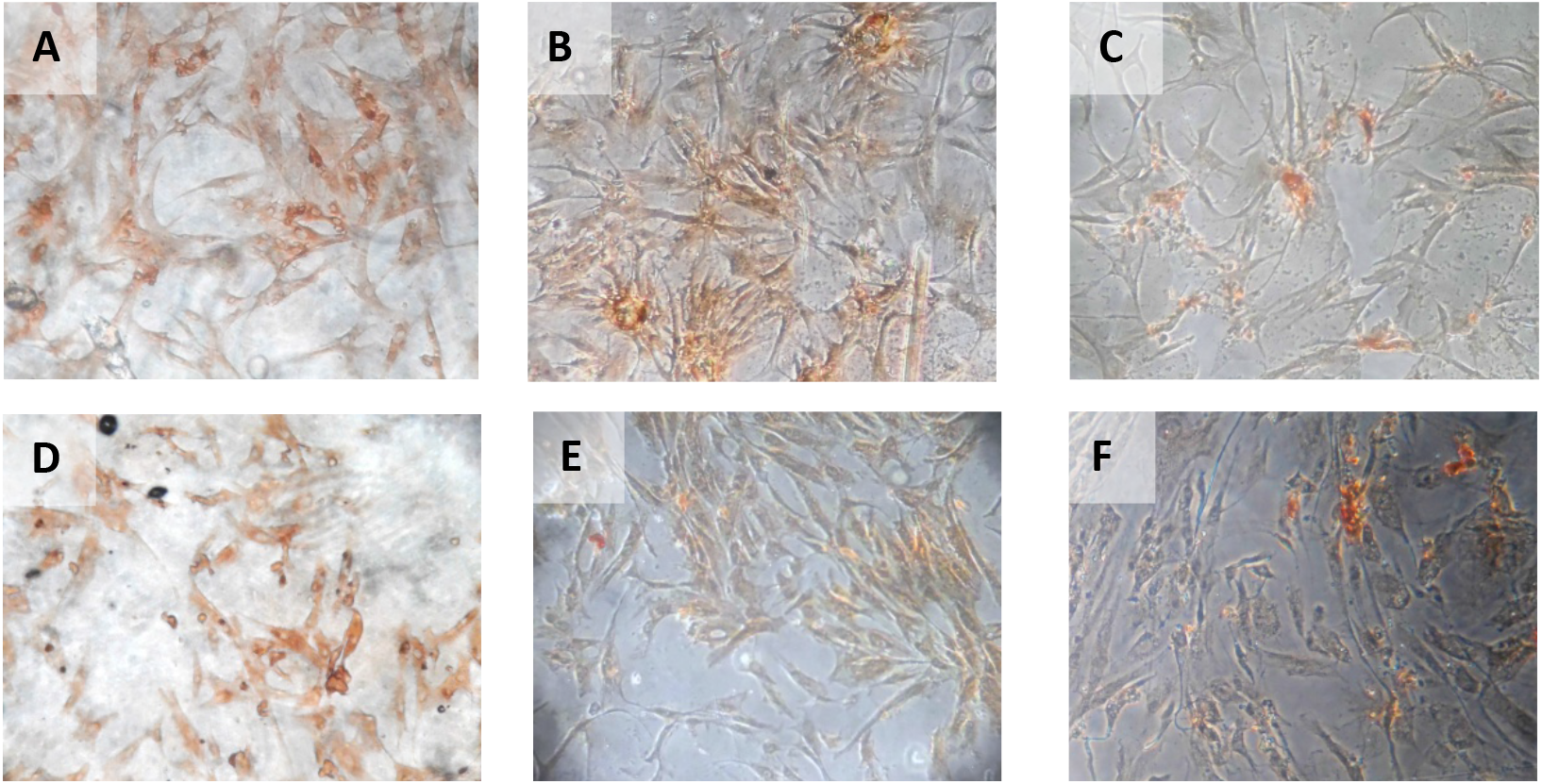
Immunohistochemical Characteristic of *h-AMSCs* based on SCF expression at: a) normoxic condition for 24 hours; b) normoxic condition for 48 hours; c) normoxic condition for 72 hours; d) hypoxic condition for 24 hours; e) hypoxic condition for 48 hours; f) hypoxic condition for 72 hours.

**Table 3.** Results on SCF expression

| Time (hour) | Experimental group | Mean ± SD | p |
| --- | --- | --- | --- |
| 24 | Normoxia | 0.084 ± 0.019 | 0.000 |
| | Hypoxia | 0.990 ± 0.013 | |
| 48 | Normoxia | 0.093 ± 0.014 | 0.000 |
| | Hypoxia | 0.901 ± 0.082 | |
| 72 | Normoxia | 0.075 ± 0.024 | 0.000 |
| | Hypoxia | 0.596 ± 0.087 | |

The result of time difference test on OCT-4 expression is between 24 hours with 48 hours (*p*<0.001), 24 hours with 72 hours (*p*<0.001), and 48 hours with 72 hours (*p*<0.001). The result of path analysis showed that SCF had an effect on OCT-4 expression (*p*<0.001; **β** =0.985). (*Table 4*) (*Figure 6*).

**Figure 6.**
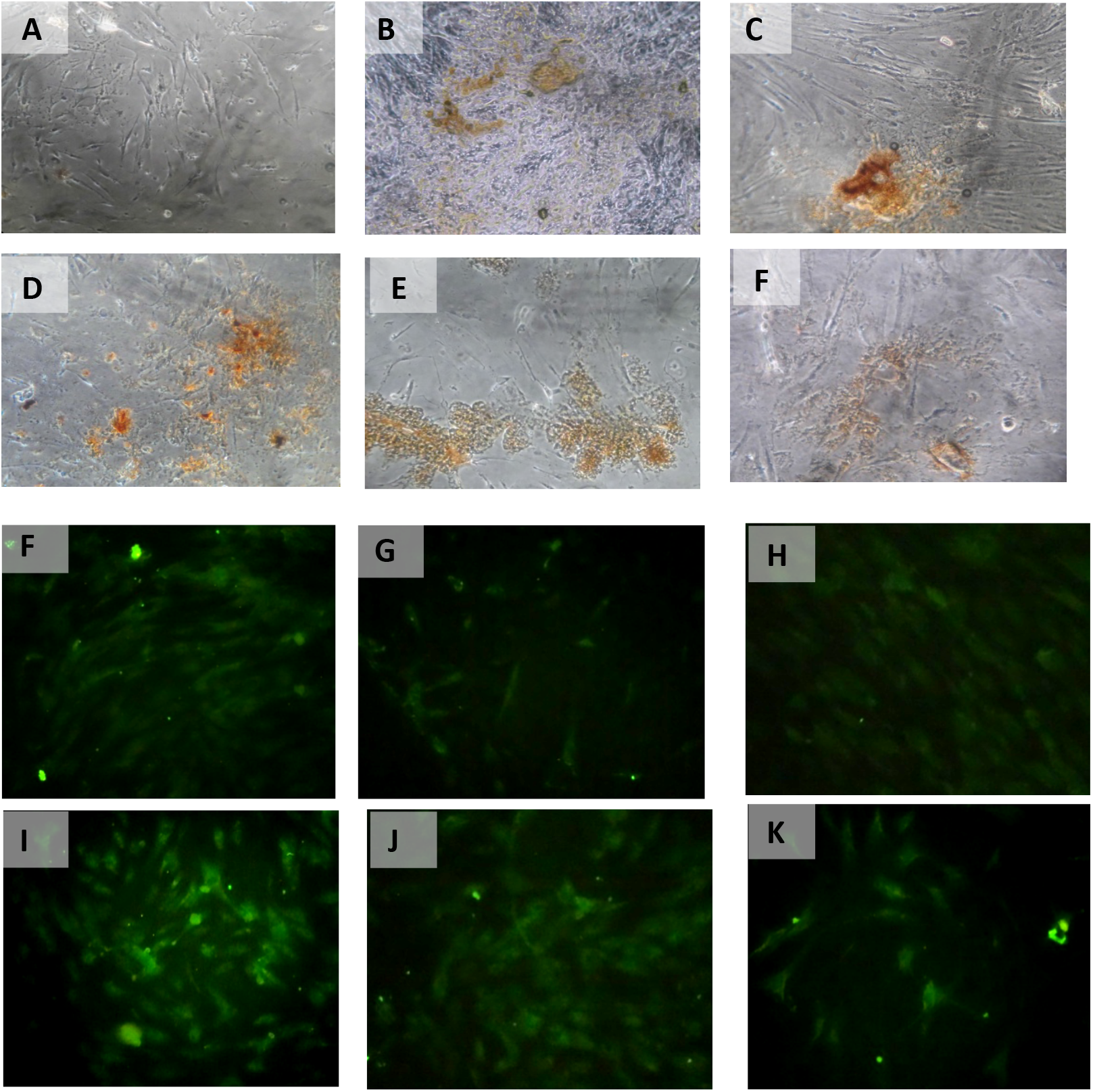
Immunohistochemical Characteristic of *h-AMSCs* based on OCT-4 expression at: a) normoxic condition for 24 hours; b) normoxic condition for 48 hours; c) normoxic condition for 72 hours; d) hypoxic condition for 24 hours; e) hypoxic condition for 48 hours; f) hypoxic condition for 72 hours. Immunofluorescence assay of *h-AMSCs* based on OCT-4 expression at: g) normoxic condition for 24 hours; h) normoxic condition for 48 hours; i) normoxic condition for 72 hours; j) hypoxic condition for 24 hours; k) hypoxic condition for 48 hours; l) hypoxic condition for 72 hours.

**Table 4.** Results on OCT4 expression

| Time (hour) | Experimental group | Mean ± SD | p |
| --- | --- | --- | --- |
| 24 | Normoxia | 0.148 ± 0.018 | 0.000 |
| | Hypoxia | 0.793 ± 0.034 | |
| 48 | Normoxia | 0.110 ± 0.007 | 0.000 |
| | Hypoxia | 0.673 ± 0.047 | |
| 72 | Normoxia | 0.099 ± 0.025 | 0.000 |
| | Hypoxia | 0.457 ± 0.151 | |

The results of time difference test on BCL2 expression between 24 hours with 48 hours (*p*=0.223), 24 hours with 72 hours (*p*=0.295), and 48 hours with 72 hours (*p*=0.982). Path analysis results show that OCT-4 effect on BCL2 expression (*p*<0.001; **β**=0.878). (*Table 5*) (*Figure 7*).

**Figure 7.**
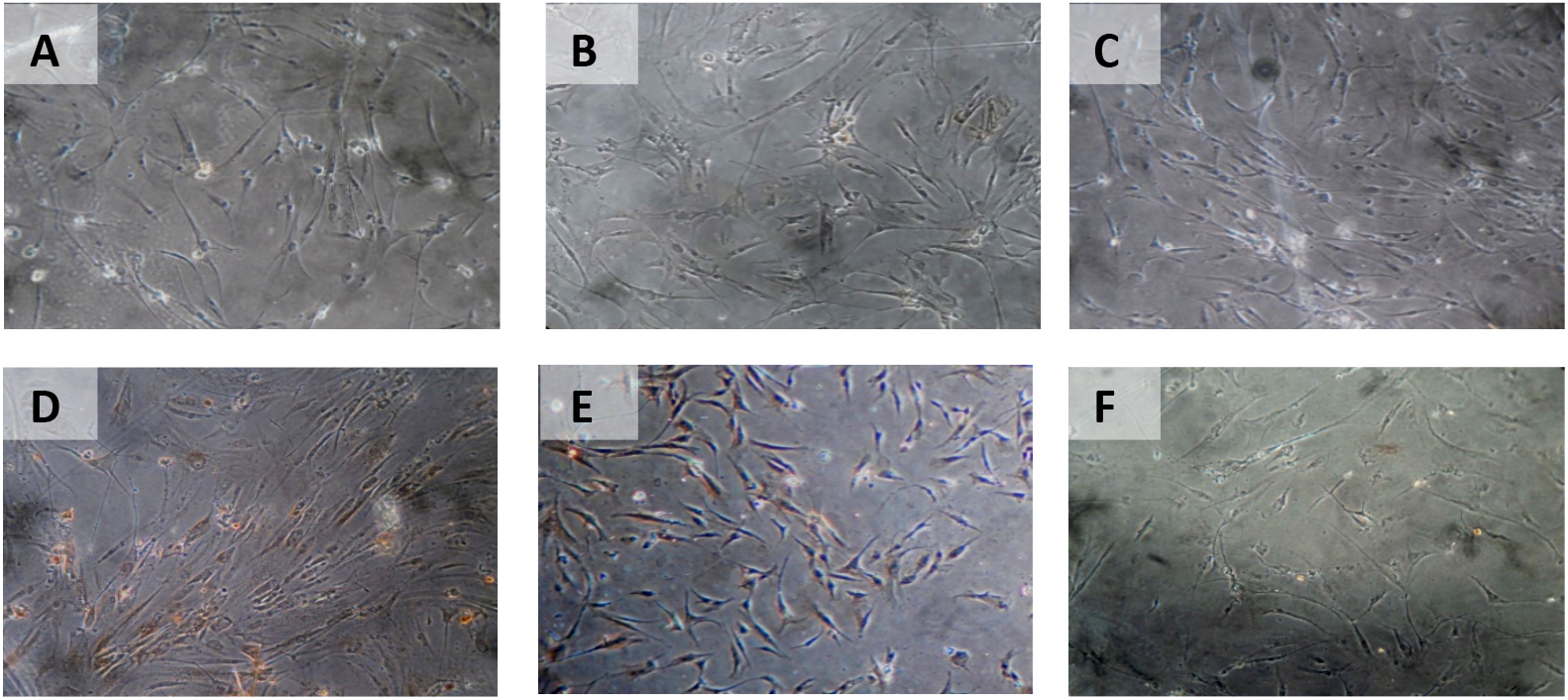
Immunohistochemical Characteristic of *h-AMSCs* based on BCL2 expression at: a) normoxic condition for 24 hours; b) normoxic condition for 48 hours; c) normoxic condition for 72 hours; d) hypoxic condition for 24 hours; e) hypoxic condition for 48 hours; f) hypoxic condition for 72 hours.

**Table 5.**
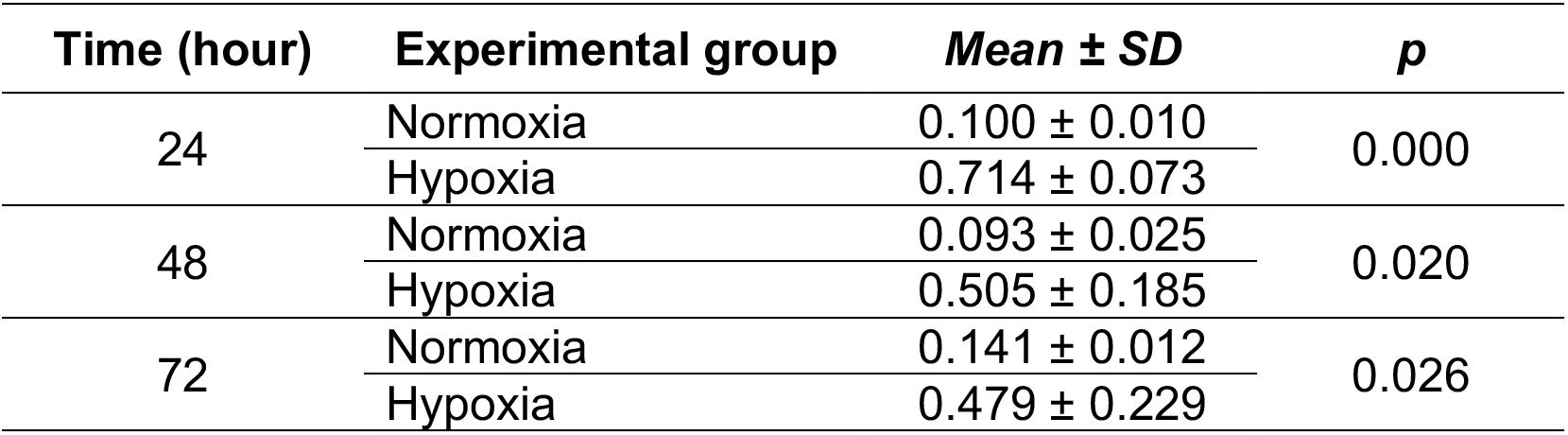
Results on BCL2 expression

The results of the time difference test on HSP27 expression between 24 hours with 48 hours (*p*=0.040), 24 hours with 72 hours (*p*<0.001), and 48 hours with 72 hours (*p*<0.001). The regression test results showed that time effects on HSP27 expression (*p*<0.001; **β** =−0.398) and hypoxia precondition also affects HSP27 expression (*p*<0.001; **β**=0.847). (*Table 6*) (*Figure 8*).

**Figure 8.**
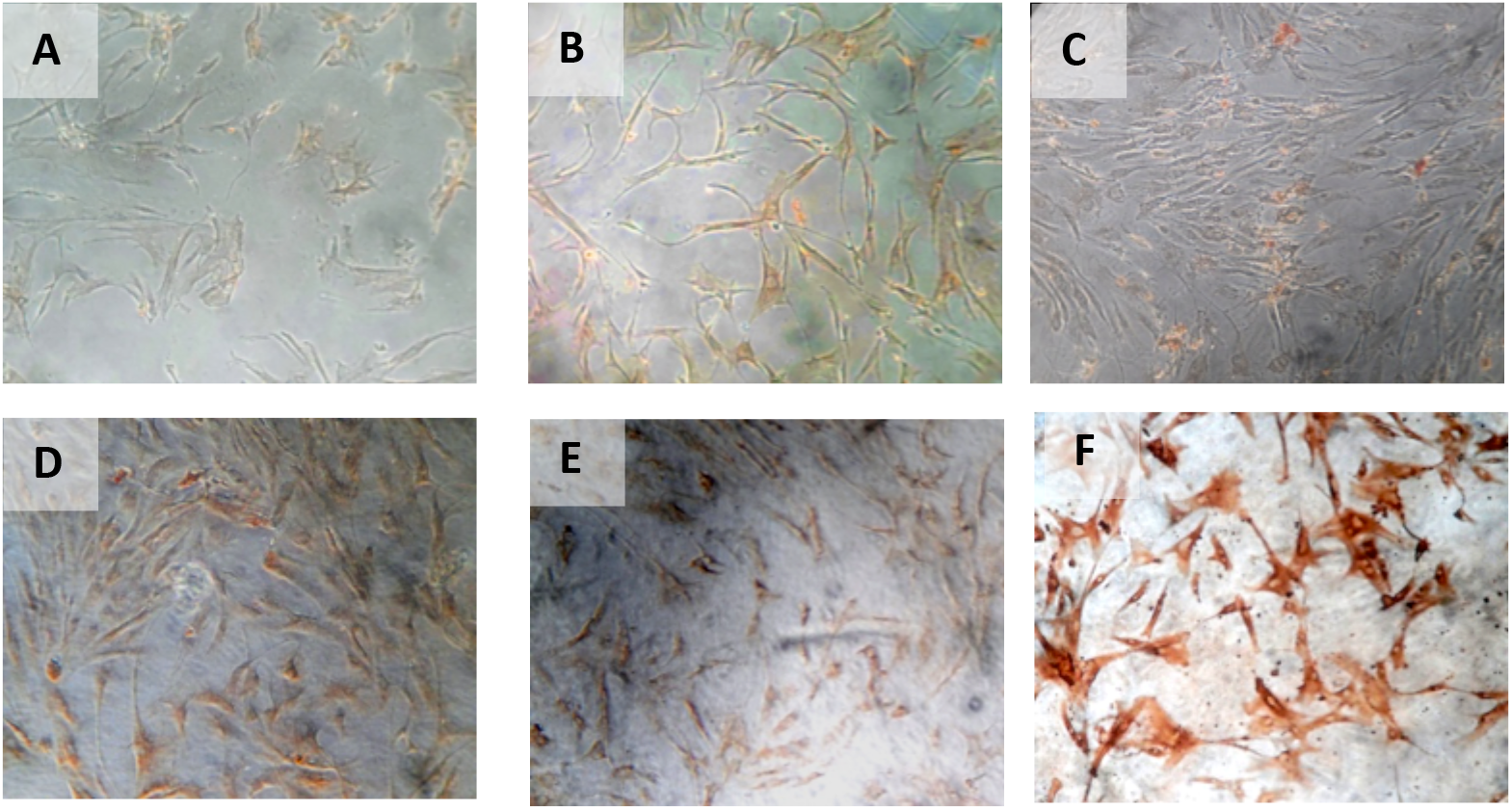
Immunohistochemical Characteristic of *h-AMSCs* based on HSP27 expression at: a) normoxic condition for 24 hours; b) normoxic condition for 48 hours; c) normoxic condition for 72 hours; d) hypoxic condition for 24 hours; e) hypoxic condition for 48 hours; f) hypoxic condition for 72 hours.

**Table 6.** Results on HSP27 expression

| Time (hour) | Experimental group | Mean ± SD | p |
| --- | --- | --- | --- |
| 24 | Normoxia | 0.156 ± 0.024 | 0.000 |
| | Hypoxia | 0.967 ± 0.018 | |
| 48 | Normoxia | 0.157 ± 0.106 | 0.000 |
| | Hypoxia | 0.773 ± 0.132 | |
| 72 | Normoxia | 0.055 ± 0.036 | 0.000 |
| | Hypoxia | 0.389 ± 0.037 | |

The results of time difference test on number of apoptotic cell amount between 24 hours with 48 hours (*p*=0.004), 24 hours with 72 hours (*p*=0.562), and 48 hours with 72 hours (*p*<0.001). Pathway analysis showed that BCL2 expression inhibited apoptosis (*p=*0.030; **β**=−0.442) and HSP27 expression also inhibited apoptosis (*p*<0.001; **β**=−0.487) (*Table 7*) (*Figure 9*).

**Figure 9.**
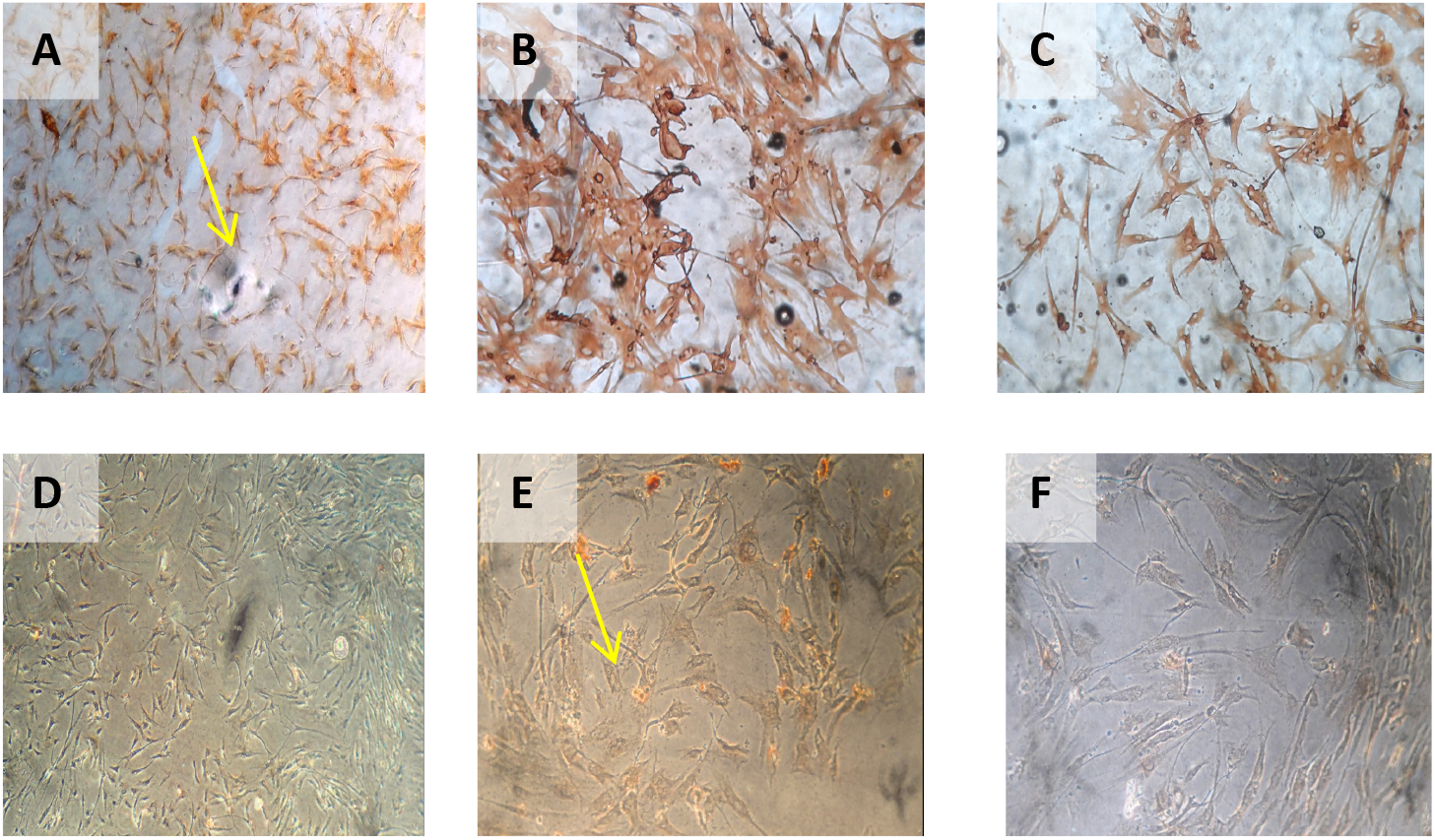
Immunohistochemical Characteristic of *h-AMSCs* based on number of apoptotic cell amount at: a) normoxic condition for 24 hours; b) normoxic condition for 48 hours; c) normoxic condition for 72 hours; d) hypoxic condition for 24 hours; e) hypoxic condition for 48 hours; f) hypoxic condition for 72 hours.

**Table 7.**
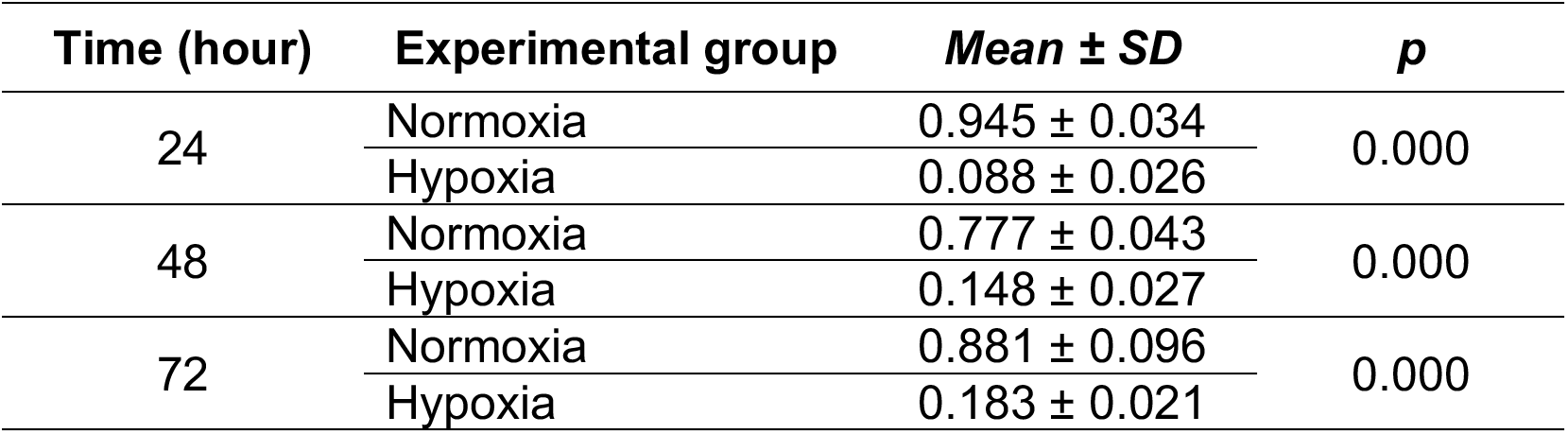
Results on number of apoptotic cell amount

| Time (hour) | Experimental group | Mean ± SD | p |
| --- | --- | --- | --- |
| 24 | Normoxia | 0.945 ± 0.034 | 0.000 |
| | Hypoxia | 0.088 ± 0.026 | |
| 48 | Normoxia | 0.777 ± 0.043 | 0.000 |
| | Hypoxia | 0.148 ± 0.027 | |
| 72 | Normoxia | 0.881 ± 0.096 | 0.000 |
| | Hypoxia | 0.183 ± 0.021 | |

## Discussion

Over the last few years, with the gradual increase in awareness of the critical role that hypoxia-induced signalling could play as a tool for generating angiogenesis on demand, two distinct approaches have emerged, as promising strategies to achieve this goal.^5^ On one hand, researchers have explored the possibility of pre-conditioning cells or grafts to hypoxia in vitro, in order to upregulate the required signalling that can then initiate angiogenesis in vivo upon transplantation.^11^ The second approach relies on direct induction of hypoxia-mediated signalling in vivo, by pharmacological means or gene therapy.^12^ A further distinction can be made on whether the therapy involves transplantation of hypoxia pre-conditioned or genetically modified cells, or if the effect is mediated directly through gene transfer or cell-free delivery of hypoxia-induced protein factors.^13^

The low survival of h-AMSCs after transplanting the heart muscle with myocardial infarction has limited the effectiveness of stem cell therapy.^8^ This is presumably because the transplanted stem cells are difficult to adapt to a new environment that is different from the environment during the in vitro culture process if it is carried out under normoxic conditions (21% oxygen concentration), while the niche of h-AMSCs in adipose tissue is actually under hypoxic conditions (oxygen concentration between 2-8%).^14^ The mechanism underlying the decreased effectiveness of stem cells when transplanted is thought to be because many transplanted stem cells undergo apoptosis.^15^ Therefore, a strategy is needed to increase the resistance of transplanted stem cells, one of which is the hypoxic adaptation process during in vitro culture.^16^

The role of sub lethal hypoxia during the in vitro culture process is to provide hypoxic preconditions so that the support niche is compatible with the hypoxic environment in vivo in myocardial infarction.^17^ Hypoxic precondition will trigger Vascular Endothelial Growth Factor (VEGF) which then binds to VEGF Receptor-1 (VEGFR-1) in the cytosol.^18^ The presence of VEGF - VEGFR-1 bonds is thought to occur in a series of signalling which activates Stem Cell Factor (SCF) or Steel Factor (SLF) in the interstitial.^19^ Interstitial SCF expression will be recognized by the SCF receptor so that an SCF-receptor complex is formed in the cell nucleus and nuclear β1-integrine expression will activate Octamer-4 (OCT-4) so that stem cells experience proliferation, self-renewal but still have the potential for differentiation.^20,21^ OCT-4 also plays a role in the activation of the PI3 / Akt pathway which affects survival cells by increasing BCL2 in the cytosol, resulting in inhibition of BAX, which causes mitochondrial PT-Pore to remain closed.^22^ The closure of the PT-Pore from the mitochondria will inhibit the release of Cytochrome-C and Apoptotic protease activating factor-1 (APAF-1) so that the apoptotic cascade does not occur.^23^

Furthermore, the hypoxic precondition will lead to the expression of Cluster of Differentiation 44+ (CD44+).^24^ This CD44+ expression occurs due to stimulation of the nuclear β1 integrin from the cell nucleus which is expressed due to the presence of the SCF-receptor complex bond.^25^ CD44+ is a hyaluronan receptor which is part of the adhesion molecule, causing interactions between cells and between cells and the matrix, as well as lymphocyte activation, also plays a role in the homing process, and increases cell migration.^26^ CD44+ is a polymorphic family that is immunologically related to proteoglycans and cell surface glycoproteins as markers of h-AMSCs. Apart from being a marker for h-AMSCs, CD44 + has a signalling function that plays a role in cell survival and motility.^26^

On the other hand, hypoxic conditions are thought to have an effect on mitochondria in increasing the expression of Reactive Oxygen Species (ROS).^27^ The increased ROS due to hypoxic conditions is thought to be the cause of the increase in free radicals formed through mitochondrial-mediated pathways.^28^ This triggers protein kinase-C (PKC) and protein K-2 (PK2) which then triggers the p53 gene so that there is an increase in p53 protein expression which will activate proapoptotic members such as BAX.^29^ Increased expression of p53 causes mitochondrial damage which causes pores to open in the membrane, so that Cytochrome-C and other molecules that act as APAF-1 will exit the mitochondria.^30^ This condition will activate procaspase 9 to become caspase-9 and followed by activation of procaspase 3 to become active caspase-3 which affects DNase so that DNA fragmentation occurs, and ends with cell death through the apoptosis process.^29^

However, the low sublethal oxygen concentration is thought to activate cells for protection in the form of repair.^31^ The repair process can be done through the activation of heat shock factor-1 (HSF-1) so that the formation of several Heat Shock Proteins (HSPs) occurs.^32^ HSPs are the product of several gene families contained in the cell nucleus which act as chaperone molecules that play a role in cell survival during the stress process.^33^ Some of the HSPs that were thought to be involved were HSP70, HSP90α and HSP27.^32^ However, in hypoxic conditions that cause the glycolysis process. This glycolysis process will further affect Krebs’s cycles so that ATP synthesis decreases.^34^ This decrease in ATP concentration is thought to cause a decrease in the function of HSP70 and HSP90α. This is because HSP70 and HSP90α are ATP-dependent chaperone molecules, thus the two HSPs (HSP70 and HSP90α) do not have the ability to act as chaperones in protecting, protecting and repairing cells under stress.^35^ The role of chaperone molecules in hypoxic conditions is carried out by HSP27, because HSP27 is ATP-independent chaperone. In addition, hypoxic precondition can maintain multipotential properties through OCT-4 expression compared to normoxic conditions.^36^

## Conclusion

From this study, it can be concluded that the hypoxic preconditioning affect the survival of h-AMSC with different apoptotic presentations due to the increased expression of BCL2 (anti apoptotic protein) and HSP 27 as chaperone proteins that play a role in inhibiting apoptosis. In this study, the hypoxic preconditioning may elevate the expression of studied variables, such as the number of apoptosis through BCL2 and HSP27 expression, trigger signal through VEGF expression, proliferation through SCF expression, and multipotency through OCT-4 expression. Hypoxic preconditioning significantly affects VEGF, VEGF affects SCF expression, SCF affects OCT-4 expression, OCT-4 affects BCL2 expression, but hypoxia also affects HSP27 expression. BCL2 and HSP27 have proven inhibiting apoptosis thus enhancing h-AMSCs survival (*Figure 10*). In conclusion, hypoxic preconditioning of h-AMSC culture has proven to increase the expression of VEGF, SCF, OCT-4, and BCL2 and HSP27. This study demonstrated and explained the existence of a new mechanism of increased h-AMSC survival in cultures with hypoxic preconditioning (O2 1%) via VEGF, SCF, OCT-4, BCL2, and HSP 27. But CD 44+ did not play a role in the mechanism of survival improvement of human AMSC survival.

**Figure 10.**
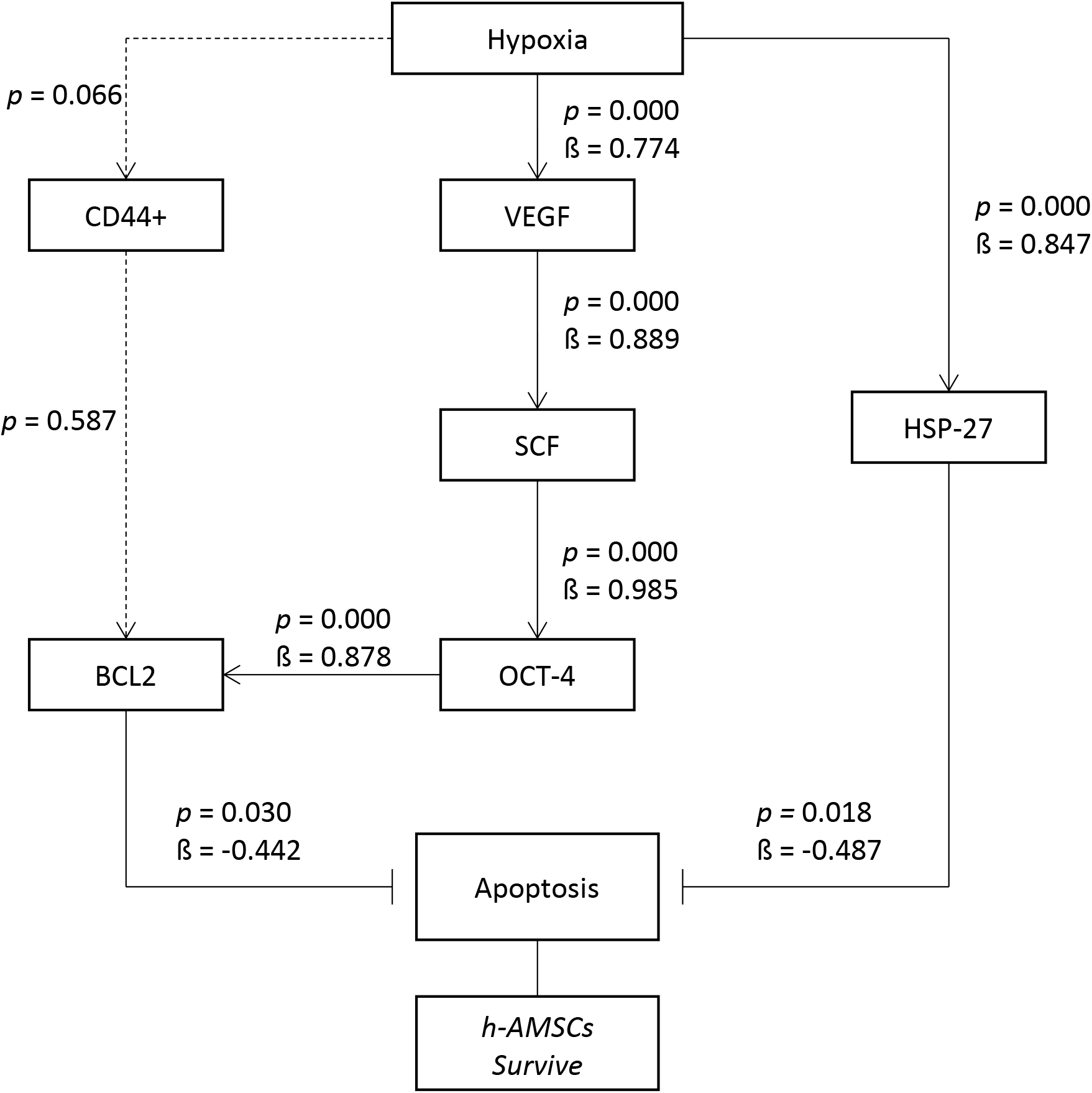
Path analysis with MANOVA and multiple linear regression analysis for hypoxic preconditioning in h-AMSCs survive

## Availability of Data and Material

The datasets generated during and/or analysed during the current study are not publicly available due to protecting participant confidentiality but are available from the corresponding author on reasonable request.

## Acknowledgement

We would also like to show our gratitude to Dr. Andrianto and Dr. Meity Ardiana for sharing their pearls of wisdom with us during the writing process, and we thank for anonymous residents and staffs for their so-called insights. We are also immensely grateful to all professors and consultants from Department of Cardiology and Vascular Medicine – Faculty of Medicine, Universitas Airlangga for their comments on an earlier version of the manuscript, although any errors are our own and should not tarnish the reputations of these esteemed persons.

## Abbreviation

AMSC: Adipose Mesenchymal Stem Cells
ATP: Adenosine Triphosphate
BAX: BCL-2-associated X protein
BCL2: B-Cell Lymphoma 2
CD44: Cluster of Differentiation 44
AMSC: human Adipose Mesenchymal Stem Cells
HSF1: Heat Shock Factor 1
HSP27: Heat Shock Protein 27
ITD: Institute of Tropical Diseases (Universitas Airlangga)
MANOVA: Multivariate Analysis of Variance
OCT4: Octamer-binding transcription factor 4
PK2: Protein k-2
PKC: Protein kinase C
ROS: Reactive oxygen species
SCF: Stem Cell Factor
SLF: Steel Factor
SPSS: Statistical Package for Social Sciences
VEGF: Vascular Endothelial Growth Factor

